## Supplemental Text for "DHX36 binding induces RNA structurome remodeling and regulates RNA abundance via m^6^A/YTHDF1"

**Inventory of Supplemental Information**

1. **Supplemental Figures**

**Figure S1.** ***In vivo* RNA structurome profiling unveils DHX36 depletion-induced global loss of RNA structures.** (A) Scatterplot showing the correlation between RT stops in two biological replicates. r represents Pearson correlation coefficient. (B) Structure-seq strategy in WT HEK293T cells was benchmarked by 18S rRNA structure at single-nucleotide resolution. 71.40% (true positive) of the bases with high SHAPE reactivity (>=0.9) correspond to single-stranded regions in the phylogenetic structure of 18S rRNA. 60.59% (true negative) of the bases with low SHAPE reactivity (<=0.6) correspond to base paired regions in the phylogenetic structure.

**Figure S2. PAR-CLIP and RBNS data analyses reveal binding preference of DHX36.** (A) The binding density of DHX36 in the designated mRNA regions. The normalized density is defined as the number of binding sites *1000 divided by the total length of the designated region. (B) The proportions of rG4-containing DHX36 binding sites in the designated regions of mRNAs. Statistical significance was calculated by Chi-square test. (C) Comparison of rG4 scores among the rG4s located in different mRNA regions. Statistical significance was calculated by unpaired two-sided Student’s t-test. (D) The distributions of R values of top 10 enriched 6mer motifs bound by RHAU53 peptides in the presence of K+ or Li+. (E) Comparison of rG4 enrichment within RHAU53-bound RNAs between K+ and Li+ conditions. Statistical significance was calculated by paired two-sided Student’s t-test. (F) The percentage of DHX36 binding sites within the designated mRNA regions covered by top10 enriched K^+^ RBNS 6mer motifs. Statistical significance was calculated by Chi-square test.

**Figure S3. DHX36 induces localized structural loss of 3’UTR binding sites.** (A) Gini index of SHAPE reactivity scores of 5’UTRs, CDSs, and 3’UTRs of DHX36-bound mRNAs in WT and DHX36-KO HEK293T cells. The fold change (FC) between KO vs. WT was shown. Statistical significance was calculated by Mann-Whitney test. (B) GC content and (C) length of the designated region of DHX36-bound mRNAs. (D) Scatterplot showing regional ΔReactivity vs. length of the designated regions of DHX36-bound mRNAs. r and P represent the correlation coefficient and statistical significance calculated by Pearson correlation test. (E) 74% of DRRs within DHX36-bound mRNAs showed decreased reactivity upon DHX36 loss. (F) The density of DRRs in the designated mRNA regions, upon normalization by DHX36 binding frequency. (G-I) The average reactivity (left) and Gini index (right) of reactivity scores of the designated mRNA region with DHX36 binding events only in (G) 5’UTR, (H) CDS and (I) 3’UTR. Statistical significance was calculated by Mann-Whitney test.

**Figure S4. DHX36 induces localized structural loss of 3’UTR binding sites.** (A) The binned average reactivity (top) and ΔReactivity (bottom) of DHX36-bound DRRs. Error bars in the bottom panel represent 95% confidence intervals (CI) of the average ΔReactivity of each bin. (B) BPP across DHX36 binding sites with and without rG4s. (C) Comparison of BPP across DHX36 binding sites located in 5’UTRs (top), CDSs (middle) and 3’UTRs (bottom). (D) The percentage of DRR-containing binding sites in the designated mRNA regions. Statistical significance was calculated by Chi-square test. (E) Localized GC content of DHX36 binding sites in the designated mRNA region. Statistical significance was calculated by Mann-Whitney test.

**Figure S5. DHX36 binding-induced structure remodeling effect is conserved in C2C12 myoblast cells.** (A) Scatterplot showing the correlation between RT stops in biological replicates 1 vs 2. r represents Pearson correlation coefficient. (B) Structure-seq strategy in WT C2C12 cells was benchmarked by 18S rRNA structure at single-nucleotide resolution. 82.80% (true positive) of the bases with high DMS reactivity (>=0.9) correspond to single-stranded regions in the phylogenetic structure of 18S rRNA. 70.10% (true negative) of the bases with low DMS reactivity (<=0.5) correspond to base paired regions in the phylogenetic structure. (C) Top: the binned average reactivity across the length of 5’UTRs (25 bins), CDSs (50 bins) and 3’UTRs (25 bins) of DHX36-bound mRNAs. Bottom: the binned ΔReactivity (KO-WT) across the above regions. Shaded area represents 95% confidence intervals of the average ΔReactivity of each bin calculated by paired two-sided Student’s t-test. (D) The average reactivity and (E) Gini index of reactivity scores of 5’UTRs, CDS, and 3’UTRs of DHX36-bound mRNAs. Mann-Whitney test was used to calculate the statistical significance. The fold change (FC) between the KO vs. WT is shown. (F) The average reactivity of DHX36 binding sites within 5’UTRs (left), CDS (middle) and 3’UTRs (right). Statistical significance was calculated by Mann-Whitney test.

**Figure S6. Transcriptomic m^6^A and YTHDF1 binding profiles.** (A) *De novo* motif discovery on m^6^A sites and YTHDF1 binding sites. (B) Venn diagram showing insignificant overlapping between pDGs with DHX36 binding to 3’UTRs and target mRNAs stabilized by YTHDF1. Hypergeometric test was used to calculate the statistical significance. (C) Comparison of average reactivity and BPP of the regions surrounding the transcriptomic m^6^A sites in WT and DHX36-KO cells. Position 0 and green area denote m^6^A residues and m^6^A motif respectively.

2. **Supplemental Tables**

Table S1. Structure-seq data analyses in HEK293T cells.

Table S2. DHX36 binding profiles in HEK293T and C2C12 cells.

Table S3. DHX36-induced DRRs within DHX36-bound mRNAs.

Table S4. Variance inflation factors (VIF) of the explanatory variables in the linear regression models.

Table S5. Structure-seq data analyses in C2C12 cells.

Table S6. Identification of DHX36 post-transcriptional regulatory targets.

Table S7. m^6^A and YTHDF1 binding sites within the DHX36 binding sites located in 3UTRs.

Table S8. Sequences of DNA, RNA oligos and peptide used in this study.
