## Supplemental Figures for "DHX36 binding induces RNA structurome remodeling and regulates RNA abundance via m^6^A/YTHDF1"

**Zhang Y and Zhao J et. al. Figure S1**

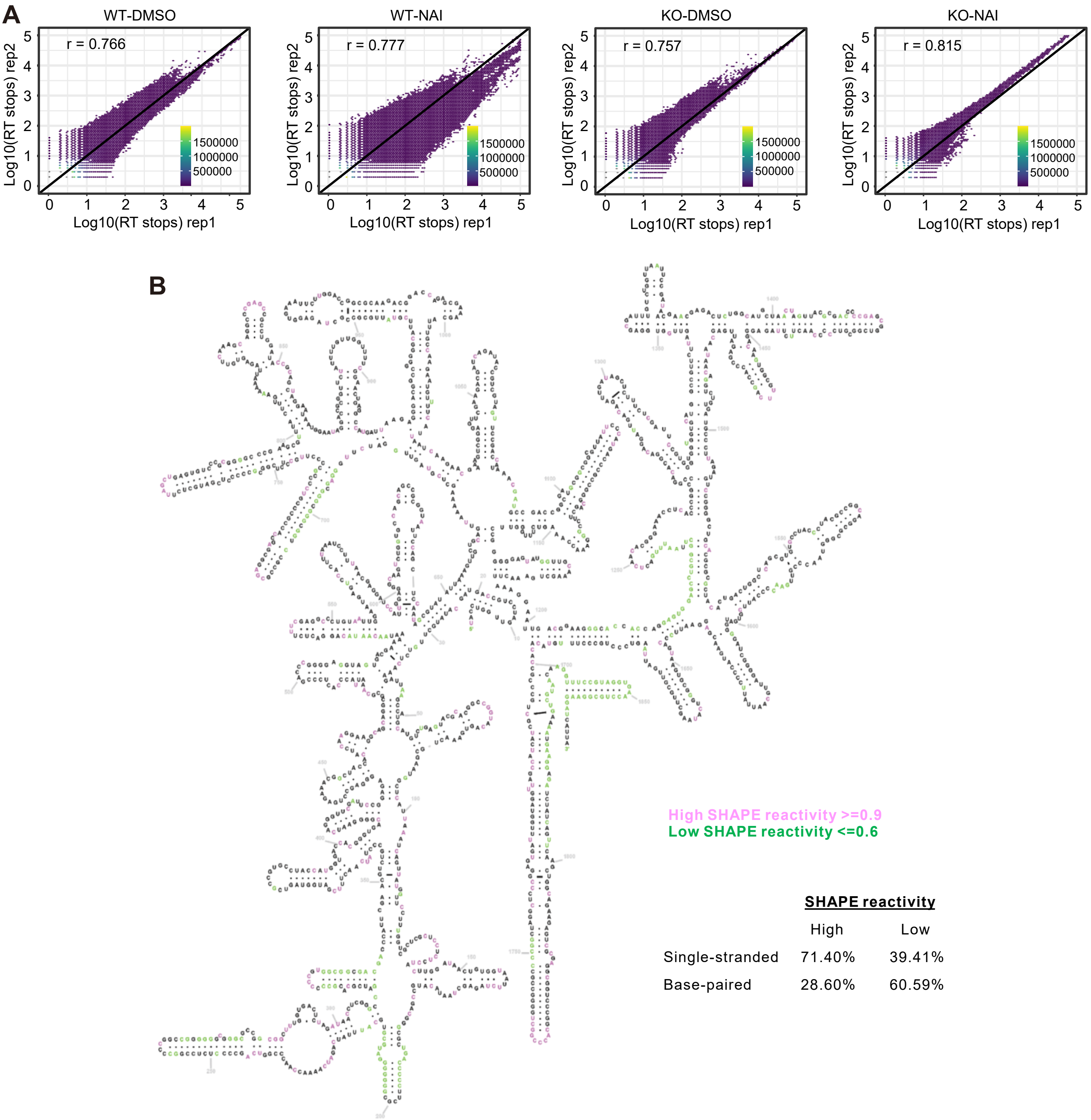


High SHAPE reactivity >=0.9
Low SHAPE reactivity <=0.6

| | SHAPE reactivity | |
|---|---|---|
| | High | Low |
| Single-stranded | 71.40% | 39.41% |
| Base-paired | 28.60% | 60.59% |

**Zhang Y and Zhao J et. al. Figure S2**

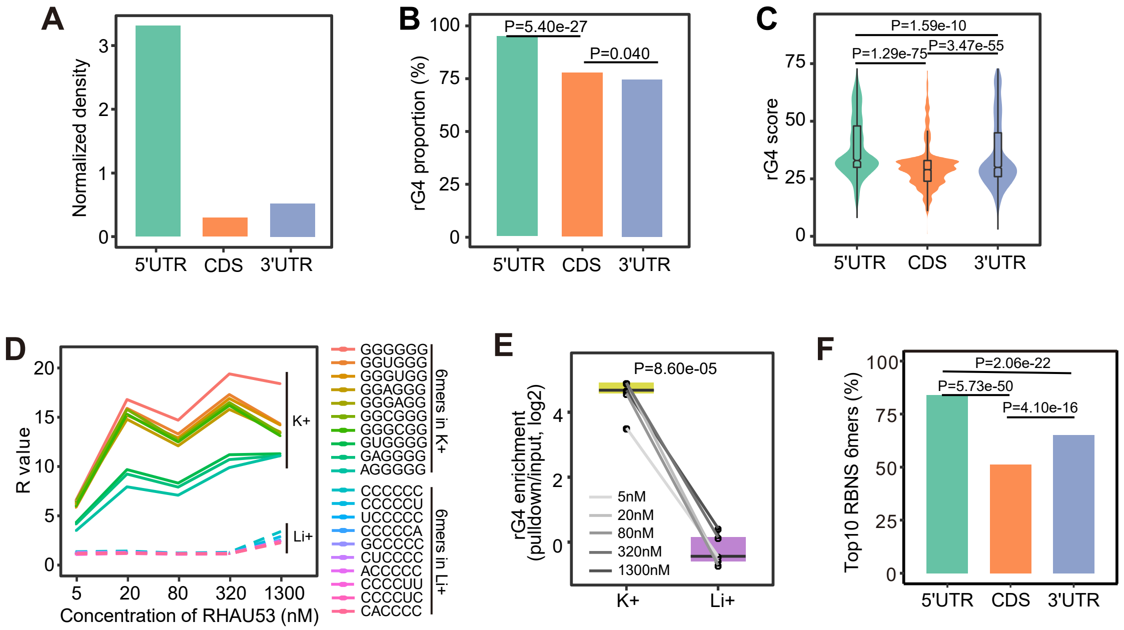

**Zhang Y and Zhao J et. al. Figure S3**

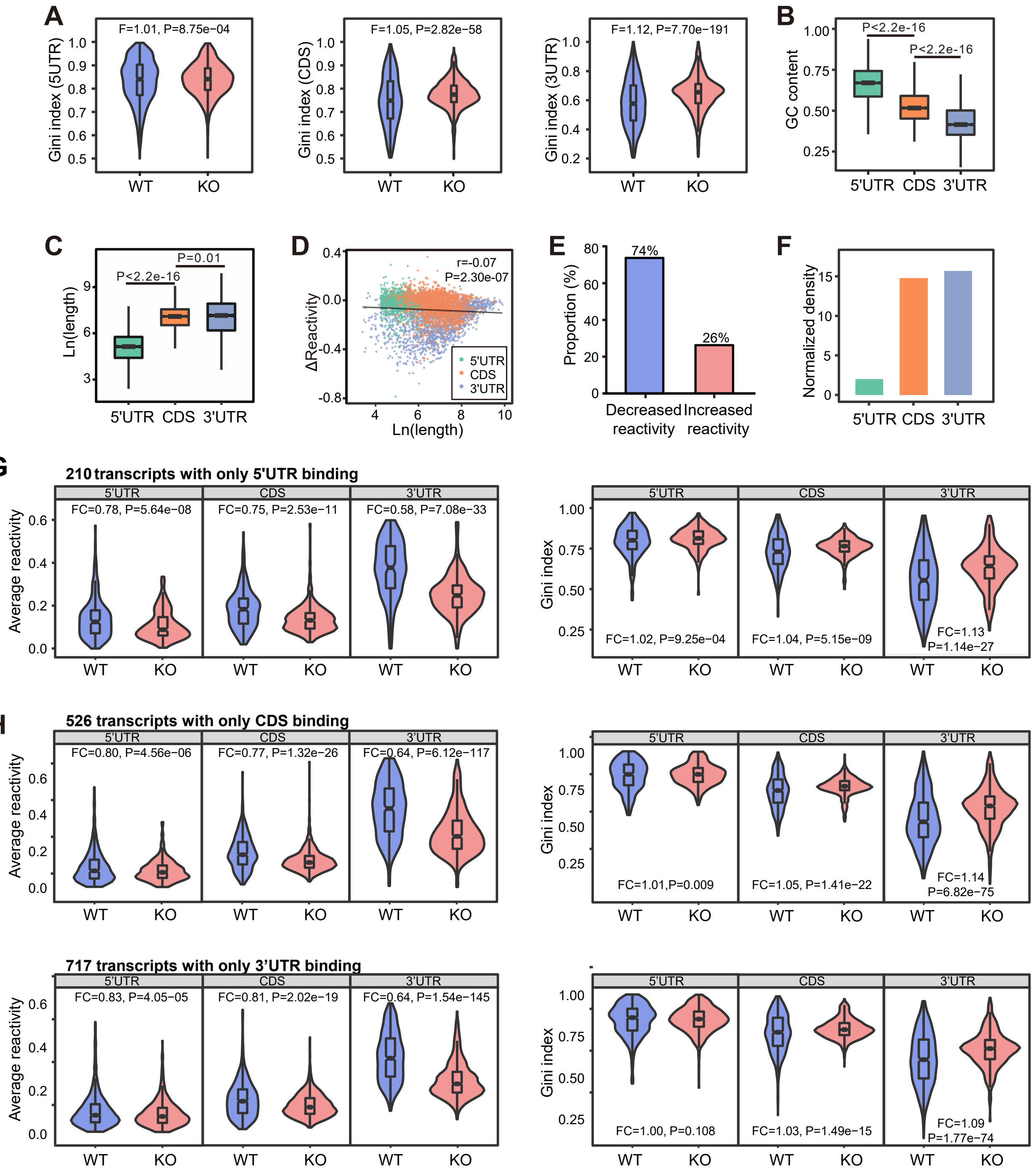

**Zhang Y and Zhao J et. al. Figure S4**

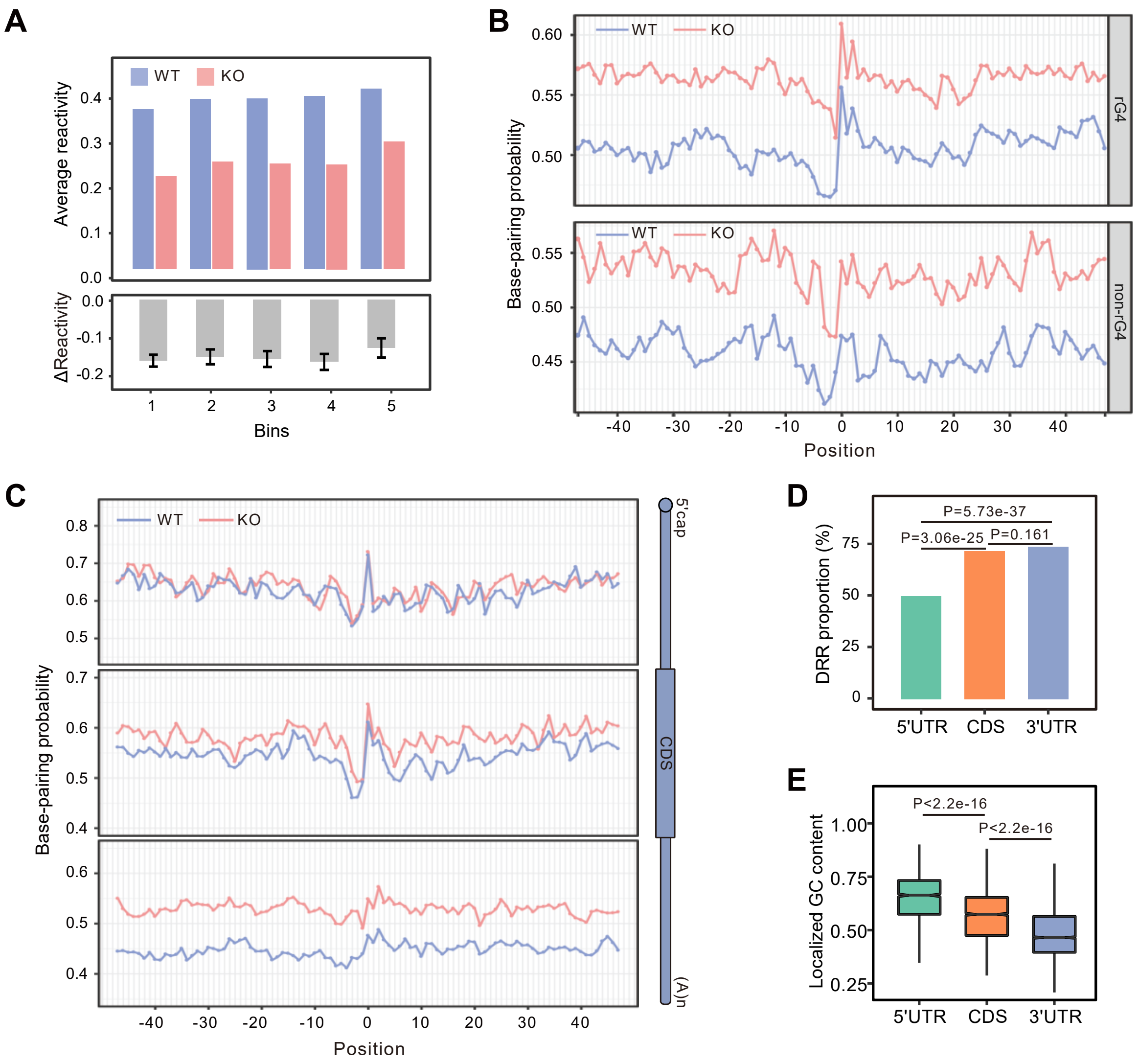

**Zhang Y and Zhao J et. al. Figure S5**

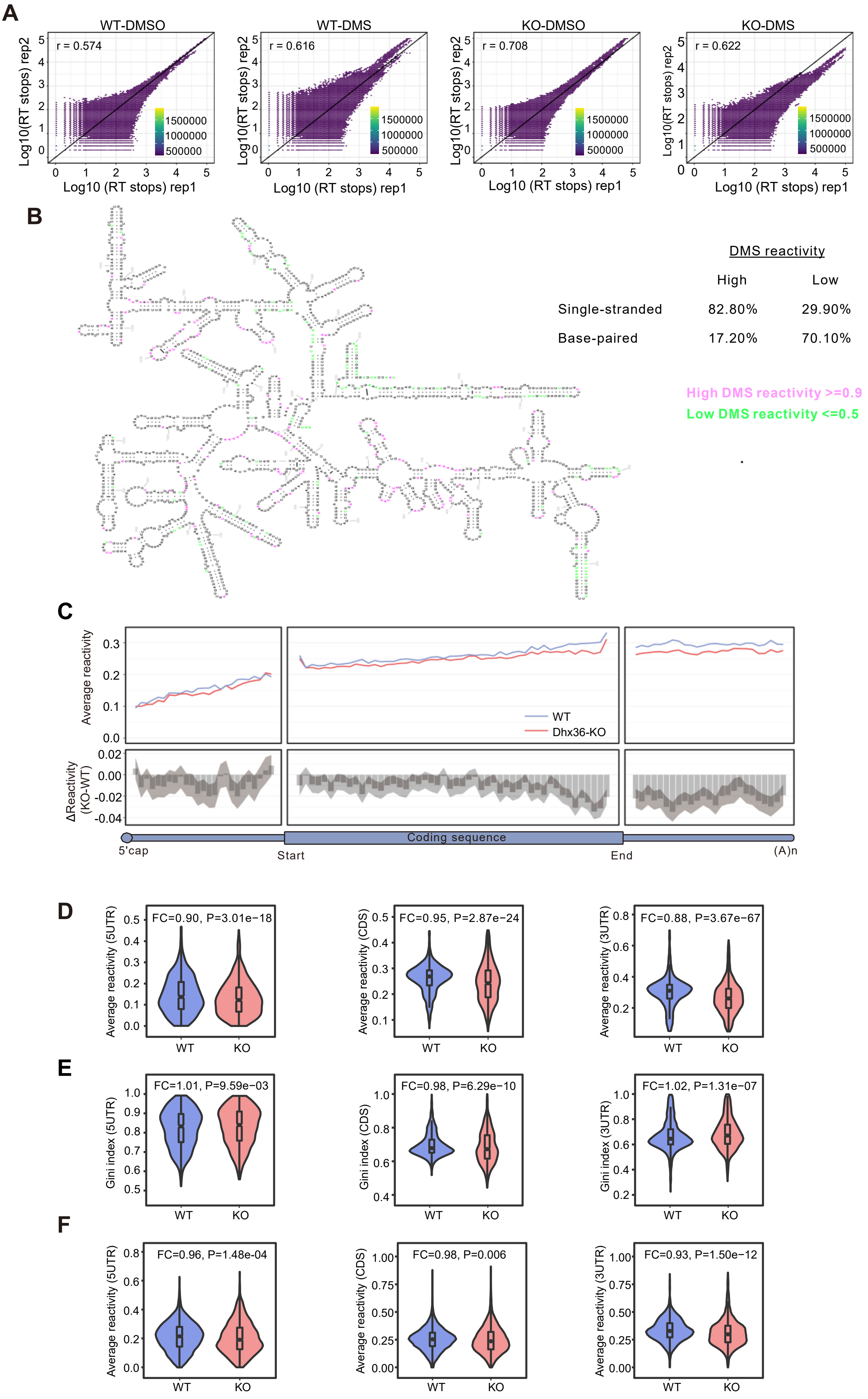

**Zhang Y and Zhao J et. al. Figure S6**

**A**

| | Motifs | P-values |
|---|---|---|
| m⁶A | 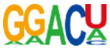 | 1e-4128 |
| YTHDF1 | 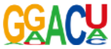 | 1e-2292 |

**B**

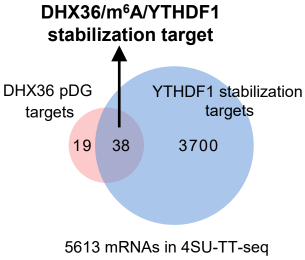


**C**

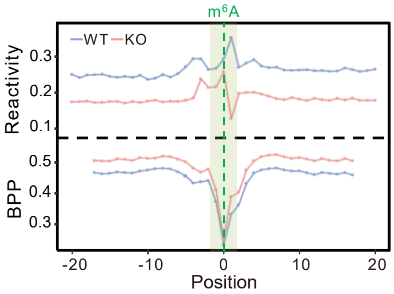
